# Intercalated amygdala dysfunction drives extinction deficits in the *Sapap3* mouse model of obsessive-compulsive disorder

**DOI:** 10.1101/2024.02.12.578709

**Authors:** Robyn St. Laurent, Kelly M Kusche, Anatol C Kreitzer, Robert C Malenka

**Affiliations:** Gladstone Institutes, San Francisco, CA, USA; Department of Physiology, Department of Neurology, Weill Institute for Neurosciences, University of California, San Francisco, CA, USA; Nancy Pritzker Laboratory, Department of Psychiatry and Behavioral Sciences, Stanford University, Stanford, CA, USA

**Author notes:** **Corresponding author:** Robert C Malenka (electronic address).

**Keywords:** amygdala, avoidance, reinforcement, extinction, obsessive-compulsive disorder, *Sapap3*

## Abstract

**Background:** The avoidance of aversive stimuli due to negative reinforcement learning is critical for survival in real-world environments, which demand dynamic responding to both positive and negative stimuli that often conflict with each other. Individuals with obsessive-compulsive disorder (OCD) commonly exhibit impaired negative reinforcement and extinction, perhaps involving deficits in amygdala functioning. An amygdala subregion of particular interest is the intercalated nuclei of the amygdala (ITC) which has been linked to negative reinforcement and extinction, with distinct clusters mediating separate aspects of behavior. This study focuses on the dorsal ITC cluster (ITC_d_) and its role in negative reinforcement during a complex behavior that models real-world dynamic decision making.

**Methods:** We investigated the impact of ITC_d_ function on negative reinforcement and extinction by applying fiber photometry measurement of GCamp6f signals and optogenetic manipulations during a platform-mediated avoidance task in a mouse model of OCD-like behavior: the *Sapap3*-null mouse.

**Results:** We find impaired neural activity in the ITC_d_ of male and female *Sapap3*-null mice to the encoding of negative stimuli during platform-mediated avoidance. *Sapap3*-null mice also exhibit deficits in extinction of avoidant behavior, which is modulated by ITC_d_ neural activity.

**Conclusions:** *Sapap3*-null mice fail to extinguish avoidant behavior in platform-mediated avoidance, due to heightened ITC_d_ activity. This deficit can be rescued by optogenetically inhibiting ITC_d_ during extinction. Together, our results provide insight into the neural mechanisms underpinning negative reinforcement deficits in the context of OCD, emphasizing the necessity of ITC_d_ in responding to negative stimuli in complex environments.

## Introduction

Appropriate selection of actions in complex environments requires behavioral flexibility, while persistent patterns of inappropriate action selection are often detrimental to well-being. Optimization of action selection is facilitated by reinforcement learning and is impaired in many neuropsychiatric disorders (1). Obsessive compulsive disorder (OCD) is one such brain disorder characterized by repetitive behaviors (i.e. compulsions), which likely develop as a form of negative reinforcement to reduce anxiety (2) and are maintained because of the inability to extinguish such learned behaviors (3–7). Thus, elucidating the circuit mechanisms mediating the extinction of negative reinforcement is one strategy for advancing our understanding of OCD pathophysiology.

A key brain area involved in positive and negative reinforcement learning is the amygdala, which is critical for deciphering the significance of positive- and negative-valence stimuli (8–10). The amygdala is comprised of distinct subregions, the functions of which are an active area of research (11). Negative reinforcement learning and extinction in a simple environment have been linked specifically to circuits involving the intercalated nuclei of the amygdala (ITC) (12–17), a conserved region also present in humans (18). The ITC consists of anatomically segregated clusters of tightly packed GABAergic neurons (19–22), which surround the basolateral amygdala (BLA). They regulate amygdala output via inhibition of the BLA and central amygdala (16, 23–26) (CeA), and reciprocal inhibition between clusters (16, 27). The distinct ITC clusters also play unique roles in regulating behavior, including the dorsal and ventromedial clusters (ITC_d_ and ITC_vm_, respectively), which have been implicated in fear learning and extinction (16).

Here, we examine the hypotheses that, (1) negative reinforcement is regulated specifically by the ITC_d_ in a dynamic reinforcement task and, (2) impaired ITC_d_ function results in negative reinforcement deficits. To better model the types of action selection for negative reinforcement required of organisms living in a dynamic environment, we used platform-mediated avoidance, a behavioral task that requires active avoidance at the cost of a positive reward (28, 29). This task is particularly appropriate for probing the role of the ITC because extinction of platform-mediated avoidance behavior has been linked to the BLA and infralimbic prelimbic cortex, regions that innervate the ITC (30). Furthermore, to investigate the relevance of extinction of negative reinforcement learning to OCD, we examined extinction of platform-mediated avoidance in the *Sapap3-*null mouse, which is commonly used to investigate behaviors related to OCD (31, 32). *Sapap3-*null mice lack the postsynaptic scaffolding protein SAP90/PSD95-associated protein 3 (*Sapap3*) (33) and display excessive grooming, increased anxiety, behavioral inflexibility, and impaired fear extinction (31, 34–38). Here, we show that ITC_d_ activity encodes threats during platform-mediated avoidance and that dysfunctional neural signals in the ITC_d_ of *Sapap3-null* mice contribute to impaired extinction of platform-mediated avoidance.

## METHODS AND MATERIALS

### Animals

Both male and female adult mice were used for all experiments and cared for in accordance with the guidelines set by the Institutional Animal Care and Use Committee at the Gladstone Institutes and Stanford University. FoxP2tm1.1(cre)Rpa/J mice (39) were originally obtained from Jackson Laboratories and the colony was maintained and bred in-house. *Sapap3* mice were originally obtained from Jackson Laboratories and heterozygotes were bred to *FoxP2*-Cre mice. We then bred pairs of *FoxP2*-cre^+^ x *Sapap3*^+/-^ mice to generate the experimental mice used for behavior. Offspring from this double transgenic cross were genotyped using Transnetyx. For all experiments, *FoxP2*-cre^+^ x *Sapap3*^+/+^ or *FoxP2*-cre^+^ x *Sapap3*^-/-^ mice were used. To reduce the potential impact of skin lesions on behavioral performance, we trimmed the hind paw nails of all mice at the time of surgery (40). For behavior experiments, mice were a minimum of postnatal day 90 at the start of the procedure. Mice were maintained on a 12-h light/dark cycle and provided food and water *ad libitum*, except where noted for water restriction in platform-mediated avoidance.

### Stereotaxic surgery

Adult male and female mice were anesthetized with 2% isoflurane gas, placed in a stereotaxic frame, and an adeno-associated virus infused through a glass microinjection pipette into the region of interest. When required, implants were slowly lowered into the brain. Optical fibers were secured to the skull with C&B Metabond and light-cured dental adhesive cement (Geristore A&B paste, DenMat). For viral injections, 100 nl of virus was infused into the ITC_d_ a rate of 50 nL/min with a borosilicate pipette coupled to a pump-mounted 5 μL Hamilton syringe (AP -0.9; ML +3.18; DV -4.6 from bregma). Pipettes remained in place for 10 min following infusion. For photometry recordings, optical fibers (Doric Lenses) with 400 μm core and 0.66 NA were unilaterally implanted over the ITC_d_ (AP -0.9; ML +3.18; DV -4.5 from bregma). For optogenetic experiments, optical fibers were made and polished using 1.25 mm diameter multimode ceramic ferrules (ThorLabs), 200 μm core fiber optic cable with 0.39 numerical aperture (NA) (ThorLabs), and 5-minute epoxy (Devcon). Experiments from mice where the injection site or implant was inaccurately placed or spread outside of the targeted region were discarded.

### Water restriction

Prior to platform-mediated avoidance, mice were placed on a water restriction regimen and slowly restricted until they achieved 85% of their original body weight. Body weight was monitored daily, before and after training sessions, and additional fluids were provided to maintain 85% body weight throughout the platform-mediated avoidance behavioral sessions.

### Platform-mediated avoidance

Training and testing is conducted in the same arena throughout the experiment: a clear, acrylic custom arena (26 x 30 x 26 cm) containing 3 custom-made infrared nose-pokes to deliver sucrose solution, a clear acrylic platform (12.5 x 12.5 cm), a shock-grid floor (MedAssociates, Inc.), and a tone generator (SparkFun WAV trigger). Acrylic arena and components are housed in a sound-attenuating chamber and illuminated with overhead LED lights. Mouse location was recorded via overhead USB camera. Task parameters and behavioral responses are triggered and recorded using custom scripts written in Statescript (Spike Gadgets) and MATLAB (Math Works) and an mbed microcontroller system connected to a computer via micro-USB. Behavior was analyzed using custom MATLAB scripts and DeepLabCut (41). This procedure is adapted slightly from previous reports using the task in rats (28, 30, 42–48). Water-restricted mice are trained for 7 sessions on a variable-interval 30s schedule where the center port pokes after the assigned ITI deliver a 10% sucrose reward to the reward port. A small light on the center port is illuminated after a correct center poke and terminates after reward retrieval. The third nose poke is inactive and triggers no outcome. On sessions 8-17, a series of 9 pairs of 30 s warning tones terminating in a 2 s (0.2 mA) footshock occur pseudo-randomly in blocks of 3 every 3 minutes. On extinction days (sessions 18-19), the warning tone occurs pseudo-randomly every 2 minutes for a total of 15 times in the absence of footshock. On recall test day (session 20), the parameters are the same as extinction days, but there are no manipulations. The sucrose schedule remains constant throughout the experiment and the clear acrylic platform is always present.

### Bulk fiber photometry imaging

Fiber photometry was performed as described previously (49). Unilateral injections of AAV1-Syn-Flex-GCaMP6f-WPRE-SV40 and implantation of a fiber optic ferrule were counterbalanced across hemispheres in separate mice. Bulk fluorescence imaging of calcium transients in freely moving mice was performed on a custom-built fiber photometry set-up. All photometry experiments were aligned to behavior using TTL pulse synchronization. Synapse software controlling an RZ5P lock-in amplifier was used to collect fiber photometry data. GCaMP6f was excited by frequency-modulated 472 and 405 nm LEDs (Doric Lenses) to stimulate Ca^+2^-dependent and isosbestic emissions, respectively. Optical signals were band-pass-filtered with a fluorescence mini cube (Doric Lenses) and signals were digitized at 6 kHz. Signal processing was performed with custom MATLAB scripts. Photometry signals were collected for each session of platform-mediated avoidance and analyzed offline by correcting the 472 nm signal for artifacts with the 405 nm isosbestic signal. Ca^+2^ signals were then z-scored for comparison across mice. Using custom-written MATLAB scripts, Ca^+2^ signals were aligned to relevant events.

### Optogenetic stimulation

Bilateral injections of AAV5-EF1α-DIO-mCherry, AAV5-EF1α-DIO-eNpHR3.0-mCherry, or AAV5-EF1α-DIO-ChR2-mCherry (titer ∼3.3-6e^12^ particles/ml) were infused into the ITC_d_. For optogenetic experiments, optical fibers were connected to a 532 nm or 473 nm laser diode (Shanghai Laser & Optics Century Co.) through a FC/PC adapter connected to a fiber optic rotary joint commutator (Doric Lenses). Statescript programs delivered continuous illumination at requisite timepoints. Fiber light output was calibrated to ∼1 mW at the fiber tip using a digital power meter console (ThorLabs).

### Histology

Mice were deeply anesthetized with ketamine (75 mg/kg) and dexmedetomidine (0.25 mg/kg) intraperitoneally and then transcardially perfused with 10 mL of phosphate-buffered saline (PBS) followed by 10 mL of 4% PFA. Whole brains were dissected out and post-fixed overnight, then transferred to 30% sucrose in PBS for 48 hours until sunk. Brains were cut to 50 μm thickness on a Vibratome Leica® 1000 plus Sectioning System and stored in PBS prior to mounting. For virus and optic fiber implant verification, slices were mounted on slides with DAPI Fluoromount-G. All brain injections and implants for behavior experiments were verified post-hoc. Slides were imaged at 4x on a Keyence slidescanner fluorescent microscope. Mistargeting of either the viral injection or implant locations were used as criteria for exclusion from behavioral analysis.

### Statistical analyses

All statistical analyses were performed in Prism (GraphPad). Where applicable, appropriate post-hoc tests were performed and p-values of p < .05 were considered significant. Statistical significance was determined using t-tests or repeated measures ANOVA (Table 1).

**Table 1.** Statistical results.

| Figure | Description | Test | Result | Post-hoc test | Result |
| --- | --- | --- | --- | --- | --- |
| 1E | Total session platform time (GCamp6f) | Repeated measures Two-way ANOVA | Interaction: $F_{1,10} = 1.993$, $p = .188$<br>Session: $F_{1,10} = 10.49$, $p = .009$<br>Genotype: $F_{1,10} = 5.326$, $p = .044$ | Šídák's multiple comparisons | WT vs. KO (Day 1) $p = .612$<br>WT vs. KO (Day 10) $p = .029$<br>Day 1 vs. 10 (WT) $p = .400$<br>Day 1 vs. 10 (KO) $p = .016$ |
| 1F | Tone platform time (GCamp6f) | Repeated measures Two-way ANOVA | Interaction: $F_{1,10} = 0.275$, $p = .612$<br>Session: $F_{1,10} = 9.284$, $p = .012$<br>Genotype: $F_{1,10} = 3.571$, $p = .088$ | Šídák's multiple comparisons | Day 1 vs. 10 (WT) $p = .199$<br>Day 1 vs. 10 (KO) $p = .059$ |
| 1G | Shocks avoided (GCamp6f) | Repeated measures Two-way ANOVA | Interaction: $F_{1,10} = 2.462$, $p = .148$<br>Session: $F_{1,10} = 10.51$, $p = .0088$<br>Genotype: $F_{1,10} = 7.50$, $p = .021$ | Šídák's multiple comparisons | WT vs. KO (Day 1) $p = .50$<br>WT vs. KO (Day 10) $p = .011$<br>Day 1 vs. 10 (WT) $p = .458$<br>Day 1 vs. 10 (KO) $p = .013$ |
| 1H | Time on platform for all avoidance sessions (GCamp6f) | Unpaired t-test, two-tailed | $t_{118} = 4.22$, $p < .0001$ | n/a | |
| 1I | Shocks avoided for all avoidance sessions (GCamp6f) | Unpaired t-test, two-tailed | $t_{118} = 5.03$, $p < .0001$ | n/a | |
| 1J | Time on platform during extinction sessions (GCamp6f) | Repeated measures Two-way ANOVA | Interaction: $F_{2,20} = 0.645$, $p = .535$<br>Session: $F_{2,20} = 5.085$, $p = .016$<br>Genotype: $F_{1,10} = 11.91$, $p = .0062$ | Šídák's multiple comparisons | Day 10 vs. Ext. Day 1 (WT) $p = .436$<br>Day 10 vs. Ext. Day 2 (WT) $p = .032$<br>Day 10 vs. Ext. Day 1 (KO) $p = .118$<br>Day 10 vs. Ext. Day 2 (KO) $p = .152$ |
| 1K | Time on platform during tone for extinction sessions (GCamp6f) | Repeated measures Two-way ANOVA | Interaction: $F_{2,20} = 2.198$, $p = .137$<br>Session: $F_{2,20} = 4.357$, $p = .027$<br>Genotype: $F_{1,10} = 6.411$, $p = .0298$ | Šídák's multiple comparisons | Day 10 vs. Ext. Day 1 (WT) $p = .578$<br>Day 10 vs. Ext. Day 2 (WT) $p = .007$<br>Day 10 vs. Ext. Day 1 (KO) $p = .508$<br>Day 10 vs. Ext. Day 2 (KO) $p = .658$ |
| 1L | Shock omission periods spent on platform for extinction sessions (GCamp6f) | Repeated measures Two-way ANOVA | Interaction: $F_{2,20} = 0.999$, $p = .386$<br>Session: $F_{2,20} = 4.059$, $p = .033$<br>Genotype: $F_{1,10} = 13.05$, $p = .005$ | Šídák's multiple comparisons | Day 10 vs. Ext. Day 1 (WT) $p = .840$<br>Day 10 vs. Ext. Day 2 (WT) $p = .049$<br>Day 10 vs. Ext. Day 1 (KO) $p = .199$<br>Day 10 vs. Ext. Day 2 (KO) $p = .235$ |
| 1M | Time on platform for all extinction sessions (GCamp6f) | Unpaired t-test, two-tailed | $t_{22} = 4.336$, $p = .0003$ | n/a | |
| 1N | "Shocks" avoided for all extinction sessions (GCamp6f) | Unpaired t-test, two-tailed | $t_{22} = 3.635$, $p = .0015$ | n/a | |
| 2F | Area under the curve (footshocks) | Repeated measures Two-way ANOVA | Interaction: $F_{1,106} = 4.636$, $p = .035$<br>Session: $F_{1,106} = 34.67$, $p < .0001$<br>Genotype: $F_{1,106} = 27.58$, $p < .0001$ | Šídák's multiple comparisons | WT vs. KO (Day 1) $p < .0001$<br>WT vs. KO (Day 10) $p = .033$<br>Day 1 vs. 10 (WT) $p < .0001$<br>Day 1 vs. 10 (KO) $p = .019$ |
| 2H | Area under the curve (extinction tones) | Repeated measures Two-way ANOVA | Interaction: $F_{1,178} = 0.017$, $p = .897$<br>Session: $F_{1,178} = 0.111$, $p = .74$<br>Genotype: $F_{1,178} = 0.155$, $p = .155$ | n/a | |
| 2J | Area under the curve (shock omissions) | Repeated measures Two-way ANOVA | Interaction: $F_{1,178} = 5.834$, $p = .017$<br>Session: $F_{1,178} = 1.59$, $p = .209$<br>Genotype: $F_{1,178} = 0.232$, $p = .631$ | Šídák's multiple comparisons | WT vs. KO (Ext. Day 1) $p = .373$<br>WT vs. KO (Ext. Day 2) $p = .096$<br>Ext. Day 1 vs. 2 (WT) $p = .658$<br>Ext. Day 1 vs. 2 (KO) $p = .02$ |
| **2L** | Peak Frequency of GCamp6f during extinction sessions | Repeated measures Two-way ANOVA | Interaction: $F_{1,10} = 1.144$, $p = .31$<br>Session: $F_{1,10} = 1.227$, $p = .294$<br>Genotype: $F_{1,10} = 6.963$, $p = .025$ | Šídák's multiple comparisons | WT vs. KO (Ext. Day 1) $p = .043$<br>WT vs. KO (Ext. Day 2) $p = .025$ |
| **2M** | Peak Amplitude of GCamp6f during extinction sessions | Repeated measures Two-way ANOVA | Interaction: $F_{1,10} = 3.700$, $p = .083$<br>Session: $F_{1,10} = 0.293$, $p = .600$<br>Genotype: $F_{1,10} = 0.731$, $p = .413$ | n/a | |
| **3F** | Time on platform during tone for early vs. late avoidance training (opto) | Repeated measures Two-way ANOVA | Interaction: $F_{1,45} = 0.139$, $p = .712$<br>Session: $F_{1,45} = 24.94$, $p < .0001$<br>Genotype: $F_{1,45} = 27.14$, $p < .0001$ | Šídák's multiple comparisons | WT vs. KO (Day 1) $p < .0001$<br>WT vs. KO (Day 10) $p = .0003$<br>Day 1 vs. 10 (WT) $p = .0003$<br>Day 1 vs. 10 (KO) $p = .008$ |
| **3H** | Shocks avoided for early vs. late avoidance training (opto) | Repeated measures Two-way ANOVA | Interaction: $F_{1,45} = 0.273$, $p = .604$<br>Session: $F_{1,45} = 27.20$, $p < .0001$<br>Condition: $F_{1,45} = 21.15$, $p < .0001$ | Šídák's multiple comparisons | WT vs. KO (Day 1) $p = .004$<br>WT vs. KO (Day 10) $p = .0004$<br>Day 1 vs. 10 (WT) $p = .0016$<br>Day 1 vs. 10 (KO) $p = .0009$ |
| **3K (WT)** | Time on platform during tone for extinction sessions (opto) | Repeated measures Two-way ANOVA (no interaction) | Session: $F_{1,15} = 0.474$, $p = .502$<br>Opto Group: $F_{1,14} = 8.113$, $p = .013$ | Šídák's multiple comparisons | mCherry vs. eNpHR3.0 $p = .035$ |
| **3K (KO)** | Time on platform during tone for extinction sessions (opto) | Repeated measures Two-way ANOVA (no interaction) | Session: $F_{1,12} = 1.17$, $p = .301$<br>Opto Group: $F_{1,11} = 9.554$, $p = .010$ | Šídák's multiple comparisons test | mCherry vs. eNpHR3.0 $p = .0046$ |
| **3L** | Time on platform during tone for recall test (opto) | Unpaired t-test, two-tailed | (WT) $t_{14} = 2.292$, $p = .038$<br>(KO) $t_{11} = 3.568$, $p = .004$ | n/a | |
| **4C (WT)** | Time on platform during tone for extinction sessions (opto) | Repeated measures Two-way ANOVA (no interaction) | Session: $F_{1,19} = 0.1856$, $p = .671$<br>Opto Group: $F_{1,18} = 0.121$, $p = .732$ | n/a | |
| **4C (KO)** | Time on platform during tone for extinction sessions (opto) | Repeated measures Two-way ANOVA (no interaction) | Session: $F_{1,13} = 0.532$, $p = .479$<br>Opto Group: $F_{1,12} = 29.49$, $p = .0002$ | Šídák's multiple comparisons test | mCherry vs. ChR2 $p < .0001$ |
| **4E** | Time on platform during tone for recall test (opto) | Unpaired t-test, two-tailed | (WT) $t_{18} = 0.151$, $p = .882$<br>(KO) $t_{12} = 2.285$, $p = .041$ | n/a | |

## RESULTS

### *Sapap3* KO mice exhibit deficits in platform-mediated avoidance

Historically*, in vivo* experiments involving manipulations of the ITC have been difficult because of the small and interspersed nature of the ITC clusters. Recently, because ITC neurons preferentially express the transcription factor, forkhead box protein P2, which is encoded by *FoxP2,* several groups have used *FoxP2*-IRES-cre mice (39) to specifically target isolated ITC clusters (16, 25, 50, 51). Because our planned experiments required transgene expression specifically in ITC_d_ neurons, all behavioral assays were performed in *FoxP2*-cre^+^ x *Sapap3^+/^*^+^ mice (“*Sapap3* WT”) or *FoxP2*-cre^+^ x *Sapap3*^-/-^ mice (“*Sapap3* KO”) (Fig 1A). These mice were subjected to platform-mediated avoidance training, which consisted of 3 phases (Fig 1B). Phase 1 (days 1-7) trained mice on a 30 second variable interval schedule where nose pokes into a center “initiation” port trigger sucrose reward from a lateral “reward” port. Reward delivery is indicated by an audible click of the solenoid used to dispense the liquid while an LED illuminated the initiation port until the reward was retrieved. During phase 2 (days 8-17), mice were presented with 9 pairs of 30 second warning tones terminating in a mild 0.2 mA footshock in blocks of 3 every 3 minutes. A safe platform was present in the corner opposite to the ports and thus the mice had to choose between avoiding the shock versus poking for sucrose, which remained available throughout the procedure. Phase 3 (days 18 and 19) extinguished avoidance behavior by providing 15 warning tones without shock at 2 min intervals.

**Figure 1.**
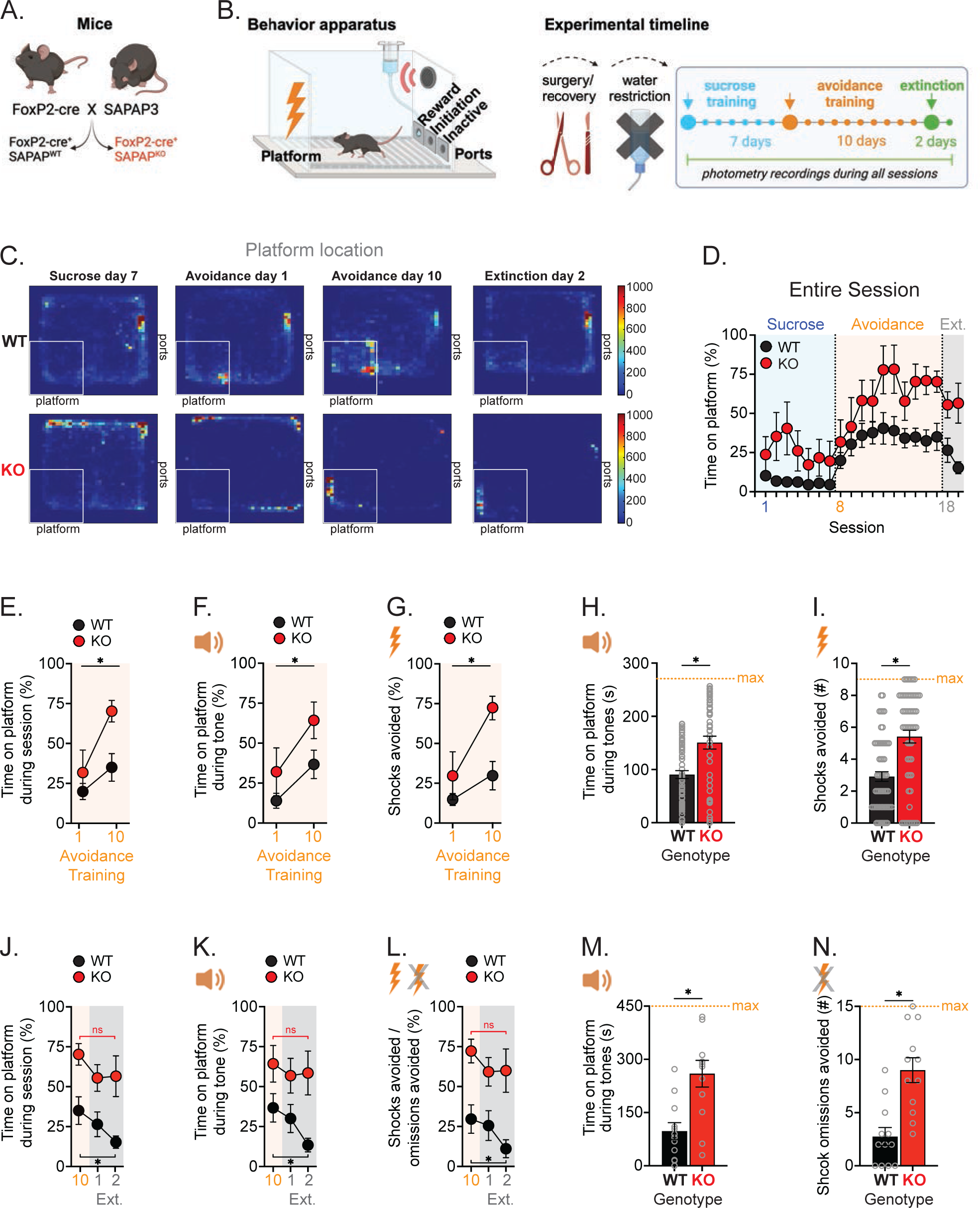
Bulk fiber photometry recording of ITC_d_ during platform mediated avoidance in *Sapap3* KO mice. *A. FoxP2*-cre and *Sapap3* mice used for experiments. *B.* Behavior apparatus and experimental timeline for platform-mediated avoidance. *C.* Heatmaps of location throughout the session for an example WT or KO mouse. *D-E.* Time on platform during entire session. *F.* Time on platform during the tone for avoidance training sessions. *G.* Percentage of shocks avoided. *H.* Time on platform during tones for all 10 avoidance training sessions. *I.* Number of shocks avoided for all 10 avoidance training sessions. *J.* Change in time on platform for the entire session from final avoidance training session through extinction. *K.* Change in time on platform during tones from final avoidance training session through extinction. *L.* Change in percentage of shock [omission] periods where the mouse was on the platform from final avoidance training session through extinction. *M.* Time on the platform during tones for both extinction sessions. *N.* Number of shock omission periods where the mouse was on the platform for both extinction sessions. n = 6 WT and 6 KO mice. Bars indicate mean ± SEM, * p < .05.

*Sapap3* KO mice exhibited several differences in the platform-mediated avoidance task with a striking difference in how much time they spent on the platform during phase 2 and 3 (Fig 1C-D). While both *Sapap3* WT and KO mice gradually spent more total time on the platform during the phase 2 avoidance sessions ([Session: F_1,10_ = 10.49, p = .009]), *Sapap3* KO mice spent significantly more time on the platform than WT mice (Fig 1E, [Genotype F_1,10_ = 5.326, p = .044]). Similarly, during the warning tone, both genotypes increased avoidance (i.e. platform time) between the first and last avoidance session with KO mice having greater tone avoidance than WT mice (Fig 1F, [Genotype F_1,10_ = 9.284, p = .012]). A complimentary metric for analyzing avoidance behavior is how many shocks the mice avoided following the warning tone. Avoided shocks increased over training sessions and was higher in *Sapap3* KO mice (Fig 1G, [Session F_1,10_ = 10.51, p = .0088, Genotype F_1,10_ = 7.50, p = .021]). Analyzing overall performance across all ten avoidance training sessions revealed that *Sapap3* KO mice spent more time on the platform during the tone (Fig 1H, [t_118_ = 4.22, p < .0001]) and avoided more shocks than *Sapap3* WT mice (Fig 1I, [t_118_ = 5.03, p < .0001]).

Importantly, during the phase 3 extinction sessions, *Sapap3* WT mice extinguished their avoidant behavior while the *Sapap3* KO mice did not. Compared to the tenth and final avoidance training session, only *Sapap3* WT mice reduced avoidance by spending less time on the platform during the session (Fig 1J, [Session F_2,20_ = 5.085, p = .016]), the tones (Fig 1K), [Session F_2,20_ = 4.357, p = .027]), and the shock omission period by the second extinction session (Fig 1L, [Session F_2,20_ = 4.059, p = .033]). Thus, *Sapap3* KO mice spent more time on the platform during the tones during extinction sessions (Fig 1M, [t_22_ = 4.336, p = .0003]). Similarly, *Sapap3* KO mice avoided more shock omission periods than *Sapap3* WT mice during extinction (Fig 1N, [t_22_ = 3.635, p = .0015]).

### Impaired ITC_d_ activity in *Sapap3* KO mice during platform-mediated avoidance

To measure neuronal population activity in the ITC_d_ during platform-mediated avoidance, we injected AAV-DIO-GCamp6f into the ITC_d_ of *Sapap3* WT or *Sapap3* KO mice (Fig 2A-B). Consistent with previous work (16, 25, 50, 51), the expression of this Cre-dependent GCamp6f was restricted specifically to the ITC_d_ (Fig 2C). On avoidance training days 1 and 10, both genotypes exhibited consistent GCamp6f responses to foot shock when located on the shock grid and diminished or absent shock responses when located on the platform (Fig 2D-E). The magnitude of these responses decreased over the course of training with smaller responses on avoidance training day 10 compared to day 1. However, ITC_d_ GCamp6f responses were smaller in the *Sapap3* KO vs. WT mice on both avoidance training days 1 and 10 (Fig 2F, [Session x Genotype *F*_1,106_ = 4.636, *p* = .035]).

**Figure 2.**
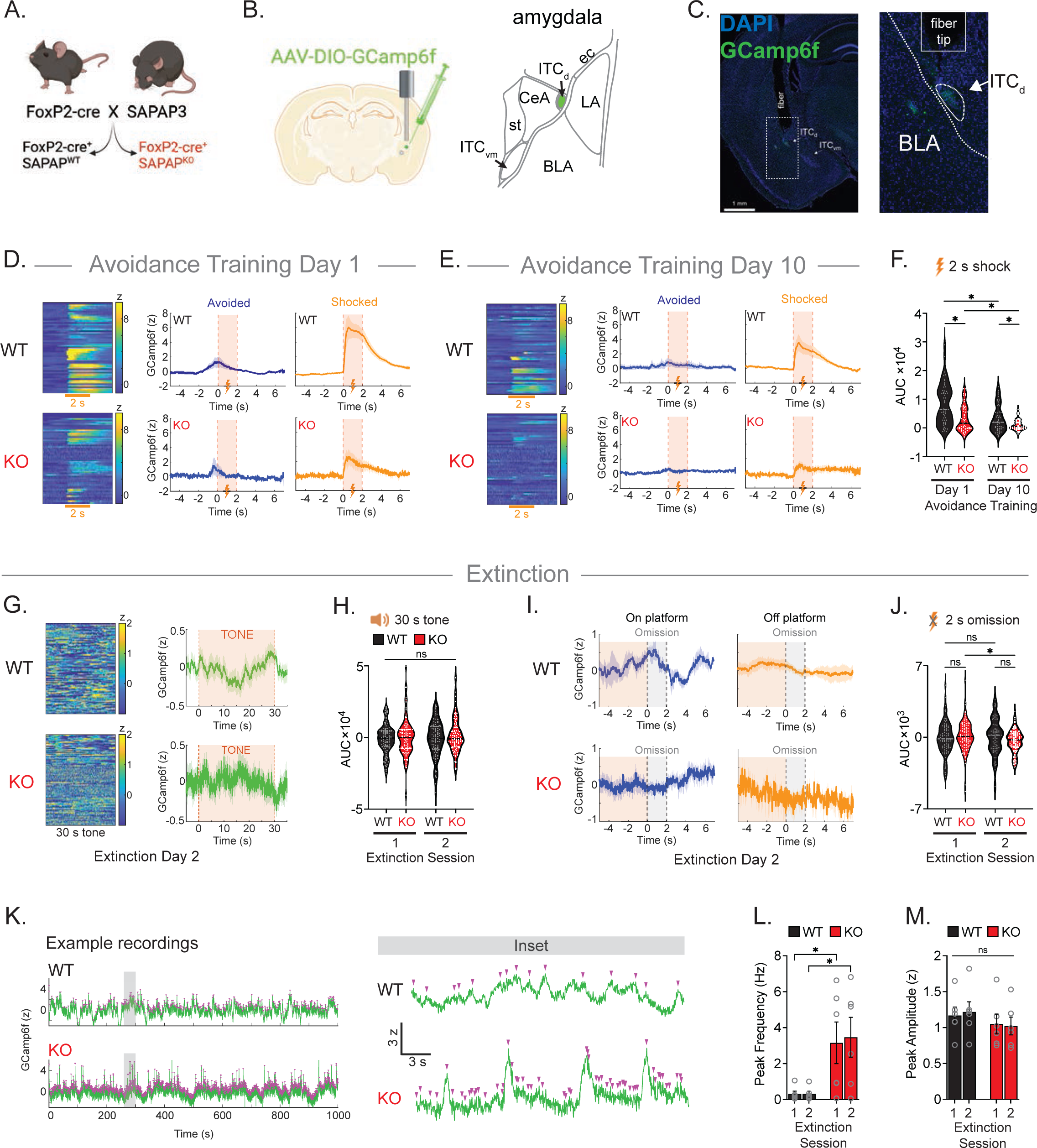
Disrupted ITC_d_ activity during platform mediated avoidance in *Sapap3* KO mice. *A. FoxP2*-cre and *Sapap3* mice used for experiments. *B.* Intracranial surgery for fiber photometry recordings. *C.* Example GCamp6f expression and fiber placement. *D.* Calcium signal aligned to 2 s shock on tone-shock avoidance training day 1 (first session). Heatmap of all trials aligned to shock (left) and Gcamp6f signal depending on the location of the mouse on or off the platform during the shock (right). n = 6 WT, 6 KO mice. *E.* Calcium signal aligned to 2 s shock on tone-shock avoidance training day 10 (final session). *F.* Area under the curve during the 2 s shock for early and late tone-shock avoidance training. Dots represent a single shock. *G.* Calcium signal aligned to 30 s warning tone during extinction day 2 for all mice, all tones. *H.* Area under the curve during the 30 s warning tone for extinction sessions. *I.* Calcium signal aligned to 2 s shock omission period and separated by location of the mouse on or off the platform during the omission period. *J.* Area under the curve during the 2 s shock omission period for extinction sessions. *K.* Example calcium signals from extinction day 2 with peak detection (left) and inset of gray bar expanded (right). *L.* Peak frequency and *M.* amplitude of calcium signal on extinction days. n = 6 WT and 6 KO mice. Abbreviations: basolateral amygdala (BLA); lateral amygdala (LA); central amygdala (CeA); stria terminalis (st); external capsule (ec); extinction (Ext.)

Arguably the most intriguing behavioral difference between *Sapap3* WT and KO mice was the failure of KO mice to extinguish their avoidance during extinction sessions (Fig 1J-N). To determine if differences in ITC_d_ activity contributed to this behavioral deficit, we examined ITC_d_ GCamp6f signals comprehensively using a variety of metrics. GCamp6f signals were not significantly different between *Sapap3* WT and KO mice during the extinction warning tones (Fig 2G-H, [Session x Genotype F_1,178_ = 0.017, p = .897]). As expected, the absence of shock following the tone resulted in reduced ITC_d_ activity (Fig 2I), similar to what was observed on avoided trials during training. Although the signal was already greatly diminished in both genotypes during the shock omission period, the GCamp6f signal modestly decreased across extinction sessions in *Sapap3* KO mice, with no change over sessions in *Sapap3* WT mice, (Fig 2J, [Session x Genotype F_1,178_ = 5.834, p = .017]). However, the most notable difference was an increase in the amount of GCamp6f transients in *Sapap3* KO mice compared to *Sapap3* WT mice (Fig 2K-L, [Genotype F_1,20_ = 6.963, p = .025]). Despite the increased rate of neural activity, the mean amplitude of GCamp6f calcium peaks were not significantly different between genotypes (Fig 2M, [(Amplitude) Genotype F_1,20_ = 0.731, p = .413]).

### Inhibition of ITC_d_ accelerates extinction

The blunted ITC_d_ GCamp6f signals in response to shock and increased spontaneous ITC_d_ GCamp6f signals during extinction of platform-mediated avoidance in *Sapap3* KO mice suggest that the ITC_d_ may play a causal role in this extinction process. To address this hypothesis, we performed optogenetic manipulations during the warning tones on extinction days in *Sapap3* WT and KO mice by injecting AAV-DIO-mCherry, AAV-DIO-eNpHR3.0-mCherry, or AAV-DIO-ChR2-mCherry bilaterally into the ITC_d_ and implanting optic fibers immediately above this structure (Fig 3A-B). The training protocol in these mice was identical to that performed in the GCamp6f-expressing mice except that we perform laser stimulation during warning tones on extinction days and added a recall test 24 h following the final extinction session to assess persistent effects of optogenetic manipulations (Fig 3C). Consistent with the results from the previous cohort of mice, *Sapap3* KO mice exhibited increased avoidance behavior (Fig 3D-H). Specifically, *Sapap3* KO mice spent more time on the platform during warning tones on the first and final avoidance training sessions compared to *Sapap3* WT mice (Fig 3F, [Genotype F_1,45_ = 27.14, p < .0001) and avoided more shocks than *Sapap3* WT mice (Fig 3H, [Genotype F_1,45_ = 21.15, p < .0001]).

**Figure 3.**
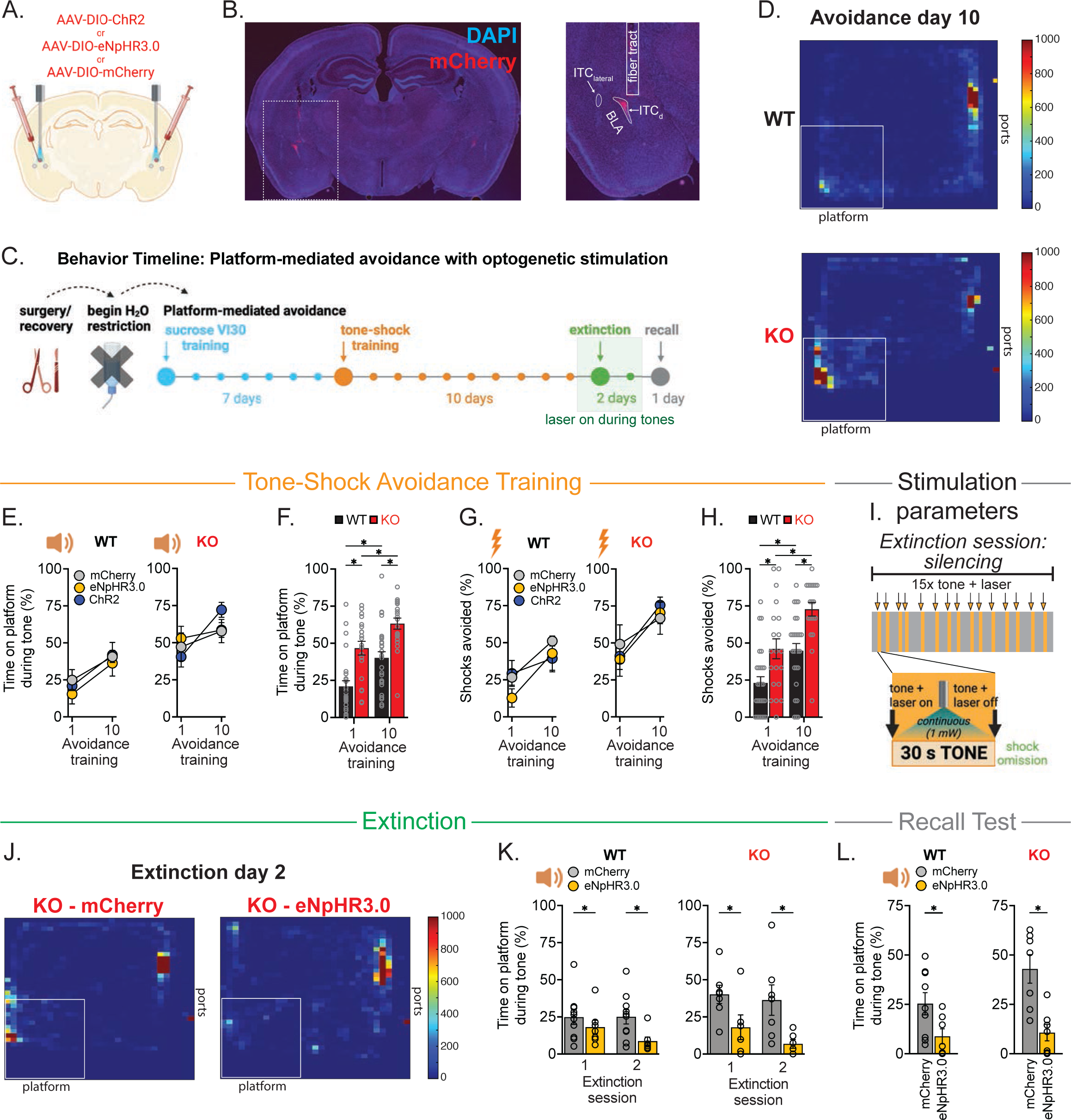
Inhibition of ITC_d_ during platform mediated avoidance enhances extinction of avoidance in *Sapap3* WT and KO mice. *A.* Intracranial surgery for bilateral viral injections and optic fiber implantation. *B.* Example viral expression and fiber placement. *C.* Experimental timeline for platform-mediated avoidance experiments. *D.* Heatmaps of location throughout the session for an example WT or KO mouse. *E-F*. Time on platform during the tone for avoidance training sessions. *G-H.* Number of shocks avoided for avoidance training sessions. *I.* Stimulation parameters for inhibition experiments during extinction sessions. *J.* Heatmaps of location for an example KO control vs. KO eNpHR3.0 mouse on extinction day 2. *K.* Time on platform during the tone for extinction sessions. *L.* Time on platform during the tone in a recall test 24 h following the final extinction session. n = 7 WT eNpHR3.0, 9 WT mCherry, 6 KO eNpHR3.0, 7 KO mCherry mice Bars indicate mean ± SEM, * p < .05.

Inhibiting ITC_d_ activity by stimulation of eNpHR3.0 during the warning tones in extinction sessions 1 and 2 enhanced extinction in both *Sapap3* WT and KO mice (Fig 3I-K). *Sapap3* WT mice expressing eNphR3.0 spent less time on the platform in the extinction sessions compared to WT mice expressing mCherry that received the same light stimulation (Fig 3K, [Opto Group F_1,14_ = 8.113, p = .013]). The same pattern occurred in *Sapap3* KO mice with the eNpHR3.0 group spending less time on the platform during tones in the second extinction session ([Opto Group F_1,11_ = 9.554, p = .01). The enhanced extinction of avoidance to the warning tone persisted into the recall test performed the following day in the absence of optogenetic manipulation for both genotypes (Fig 3L, [WT: t_14_ = 2.292, p = .038], [KO: t_11_ = 3.568, p = .004]).

### Activation of ITC_d_ enhances extinction deficit in *Sapap3* KO mice

Given that inhibition of ITC_d_ activity during the warning tones enhanced extinction, we next addressed whether increasing ITC_d_ activity during warning tones (Fig 4A) does the opposite, as evidenced by increased time spent on the platform during extinction sessions. Optogenetic activation of ChR2 in ITC_d_ in *Sapap3* KO mice dramatically increased time spent on the platform during the warning tone in both extinction sessions compared to mCherry expressing mice (Fig 4B-C, [Opto Group F_1,12_ = 29.49, p = .0002]). Surprisingly, optogenetic stimulation of ITC_d_ neurons in *Sapap3* WT mice had no effect on time spent on the platform during the warning tones compared to mCherry controls ([Opto Group F_1,18_ = 0.121, p = .732]). This manipulation in *Sapap3* WT mice also did not influence performance in the recall test following extinction (Fig 4D-E, [WT: t_18_ = 0.151, p = .882]). In contrast, in *Sapap3* KO mice, the failure to extinguish avoidance to the warning tone persisted into the recall test in the absence of optogenetic stimulation ([KO: t_12_ = 2.285, p = .041]).

**Figure 4.**
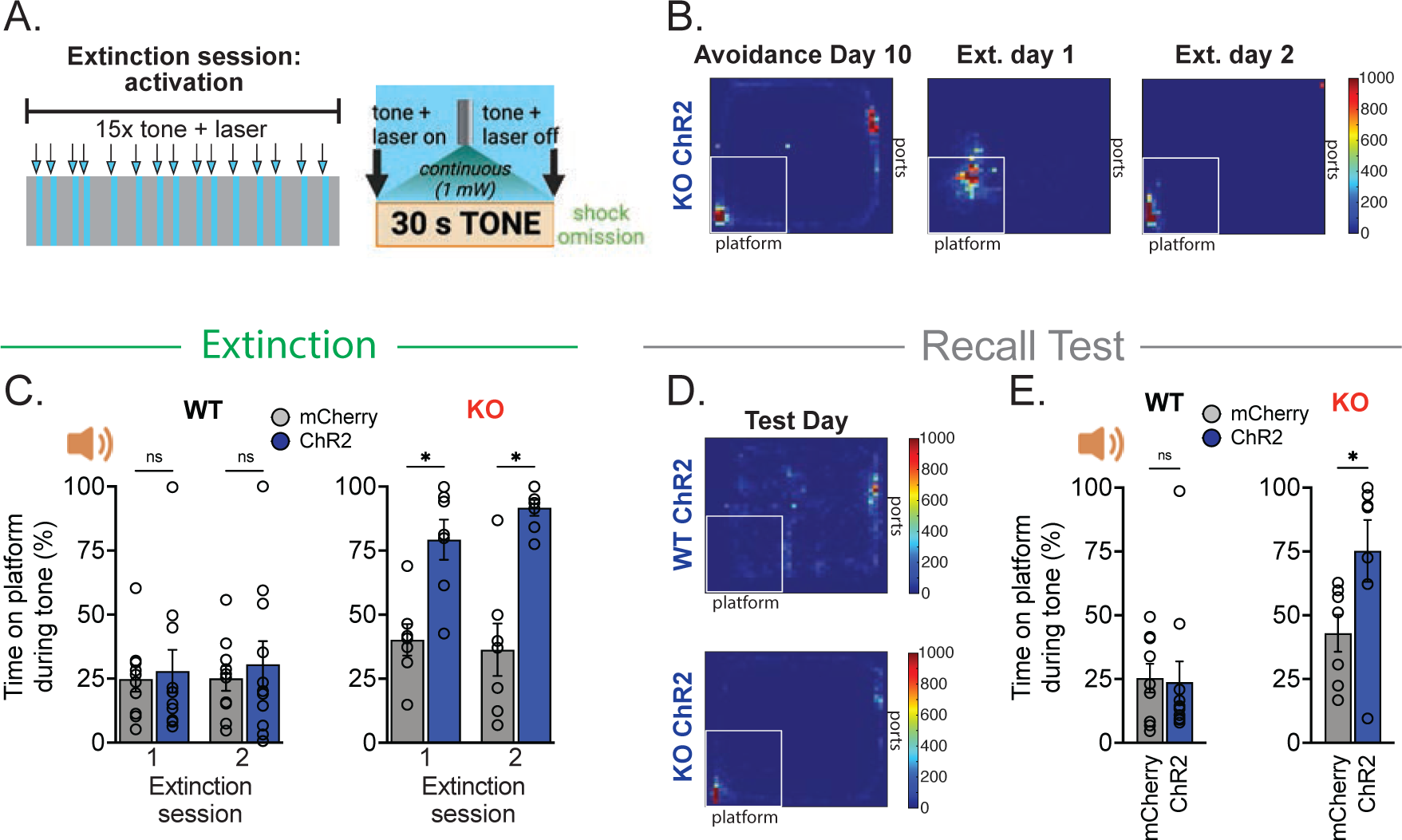
Activation of ITC_d_ during platform mediated avoidance impairs extinction of avoidance in *Sapap3* KO mice. *A.* Stimulation parameters for silencing experiments during extinction sessions. *B.* Heatmaps of location throughout the session for an example KO mouse. *C*. Time on platform during the tone for extinction sessions. *D*. Heatmaps of location for an example WT ChR2 vs. KO ChR2 mouse on recall test day. *E*. Time on platform during the tone in a recall test 24 h following the final extinction session. n = 11 WT ChR2, 9 WT mCherry, 7 KO ChR2, 7 KO mCherry mice Bars indicate mean ± SEM, * p < .05.

## Discussion

### Summary

Here, we compare the function of the ITC_d_ in *Sapap3* WT and KO mice using a platform-mediated avoidance task involving competing motivations for positive and negative stimuli. *Sapap3* KO mice spent more time on the platform during avoidance training and, most importantly, unlike WT mice, failed to extinguish this avoidance behavior. These aberrant behaviors in the *Sapap3* KO mice were associated with a reduction in the ITC_d_ neural response to a footshock as well as an increase in spontaneous Ca^+2^ transients during extinction indicating heightened ITC_d_ activity. Consistent with a causal role for increased ITC_d_ activity in the extinction deficit in *Sapap3* KO mice, optogenetic inhibition of ITC_d_ neurons rescued extinction of the avoidant behavior. The same manipulation also facilitated extinction in *Sapap3* WT mice. In contrast, optogenetic activation of the ITC_d_ resulted in complete impairment of extinction in *Sapap3* KO mice with no effect on WT mice. Importantly, these interventions provided lasting efficacy, with the modification of extinction persisting into a recall test 24 h after the optogenetic manipulations.

### Role of the ITC in negative reinforcement learning

Reinforcement learning allows for optimization of behavior selection and is mediated in part by brain circuits involving the striatum and amygdala (52). The ITC has been suggested to play a particularly important role in negative reinforcement and extinction (12–17), which are processes impaired in OCD subjects (4–7, 53). Here, our goal was to understand the potential role of a single ITC cluster, the ITC_d_, in encoding negative reinforcement and extinction during a complex approach-avoidance task. Much of our understanding of the role of the ITC in negative reinforcement has been limited by the inability to target individual ITC clusters. Most prior studies used classic fear conditioning which measures a passive response to a threat, with the exception of a correlative study showing increased neural activation after extinction of lever-pressing to avoid a shock (14). Using fear conditioning, previous reports showed that partial lesions of the ITC led to impaired fear extinction (12) and distinct ITC clusters exhibited neural activation after fear learning or extinction (13). ITC clusters might regulate the switch between fear and extinction behaviors via changes in synaptic strength, for instance, BLA inputs to the ITC are altered with fear extinction (54–56). A recent report using *FoxP2*-cre mice found that ITC_d_ neurons encode fear memory and conversely, ITC_vm_ neurons encode extinction (16). Furthermore, inactivation of ITC_d_ reduced freezing and inactivation of ITC_vm_ increased freezing during fear memory retrieval (16). Our data are in line with these findings, suggesting that ITC_d_ encoding of responses to negative stimuli in simple environments extends to complex operant tasks, such as the platform-mediated avoidance task used here.

### Effects of Sapap3 deletion

*Sapap3* KO mice have been extensively investigated as a rodent model of OCD because they express repetitive and compulsive behaviors, reinforcement deficits, increased anxiety, as well as improvement in symptom severity with serotonin reuptake inhibitor treatment (31, 34, 36–38, 57–60). Several of the *Sapap3* KO behavioral abnormalities appear to be due to deficits in cortico-striatal synaptic transmission (31, 61, 62), consistent with the generally accepted hypothesis that OCD involves abnormal functioning of cortico-striatal-thalamic-cortical networks (63). In addition to its high expression in the striatum, *Sapap3* is expressed in many regions of the amygdala in rodents (31, 33, 64). Although further work will be necessary to delineate how *Sapap3* deletion causes deficits in ITC_d_ activity, given its prominent role in maintaining excitatory synaptic function, it seems likely that a reduction in synaptic transmission at glutamatergic synapses onto ITC_d_ neurons is one likely contributing mechanism. Another possibility is changes in synaptic properties or intrinsic excitability of ITC_d_ neurons due to homeostatic plasticity because of the loss of *Sapap3* throughout development.

We obtained two surprising results when manipulating ITC_d_ during extinction. First, the optogenetic manipulations that were applied only during the two extinction sessions caused behavioral changes that were still present 24 hours later. This suggests that suppression of ITC_d_ activity during extinction allows for generation of more robust extinction memory, perhaps via removal of inhibition onto ITC_vm_ extinction neurons through the reciprocal connection of ITC_d_→ITC_vm_. Second, optogenetic activation of ITC_d_ in *Sapap3* KO mice exacerbated their extinction deficits yet had no effects on extinction of platform-mediated avoidance in *Sapap3* WT mice. Similar to human OCD subjects (37), this difference could be explained by *Sapap3* KO mice responding to negative stimuli more robustly. Additionally, deficits in ITC_vm_ or its inputs could lead to an imbalance between the reciprocally connected ITC_d_ and ITC_vm_ in *Sapap3* KO mice, allowing for our moderate activation with ChR2 to overpower the inhibition originating in the ITC_vm_. Conversely, in WT mice, endogenous ITC_vm_→ITC_d_ inhibition may be sufficient to override our optogenetic manipulations in WT but not KO mice.

### Conclusions

Morphological and functional abnormalities in the amygdala of OCD subjects have been reported (65–73) but little is known about the potential role of amygdala dysfunction in contributing to negative reinforcement deficits in OCD. Using a mouse model for OCD, we present evidence that abnormalities in the functioning of a specific subregion of the amygdala, the ITC_d_, importantly contributes to impairment in the extinction of an avoidant behavior. Specifically, we identified diminished neural responses in the ITC_d_ to an aversive stimulus and increased neural activity during extinction. Most importantly, we demonstrate that inhibition of ITC_d_ accelerates the extinction process and persistently rescued the extinction deficit in *Sapap3* KO mice, suggesting that targeting ITC_d_ circuits in patients with impaired negative reinforcement deficits has therapeutic potential.

## ACKNOWLEDGEMENTS

This work was supported by a grant from the Foundation for OCD Research (AK, RM) and the Berkelhammer Award for Excellence in Neuroscience (RS). The authors would like to thank Kendall Raymond for assisting in imaging and Aphroditi Mamilagas for assistance with MATLAB scripts to analyze calcium imaging peak frequency & amplitude. Some schematics included in figures were created with BioRender.

CRediT author statements for this manuscript: Robyn St. Laurent participated in Conceptualization, Methodology, Software, Validation, Formal Analysis, Investigation, Resources, Data Curation, Writing – Original Draft, Writing – Review & Editing, Visualization, and Funding Acquisition. Kelly Kusche participated in Investigation and Writing – Review & Editing. Anatol Kreitzer participated in Conceptualization, Writing – Review & Editing, Supervision, Project Administration, and Funding Acquisition. Robert Malenka participated in Conceptualization, Writing – Review & Editing, Supervision, Project Administration, and Funding Acquisition.

Data and code used for statistical analyses presented here are available upon request to the corresponding author.

## KEY RESOURCES TABLE

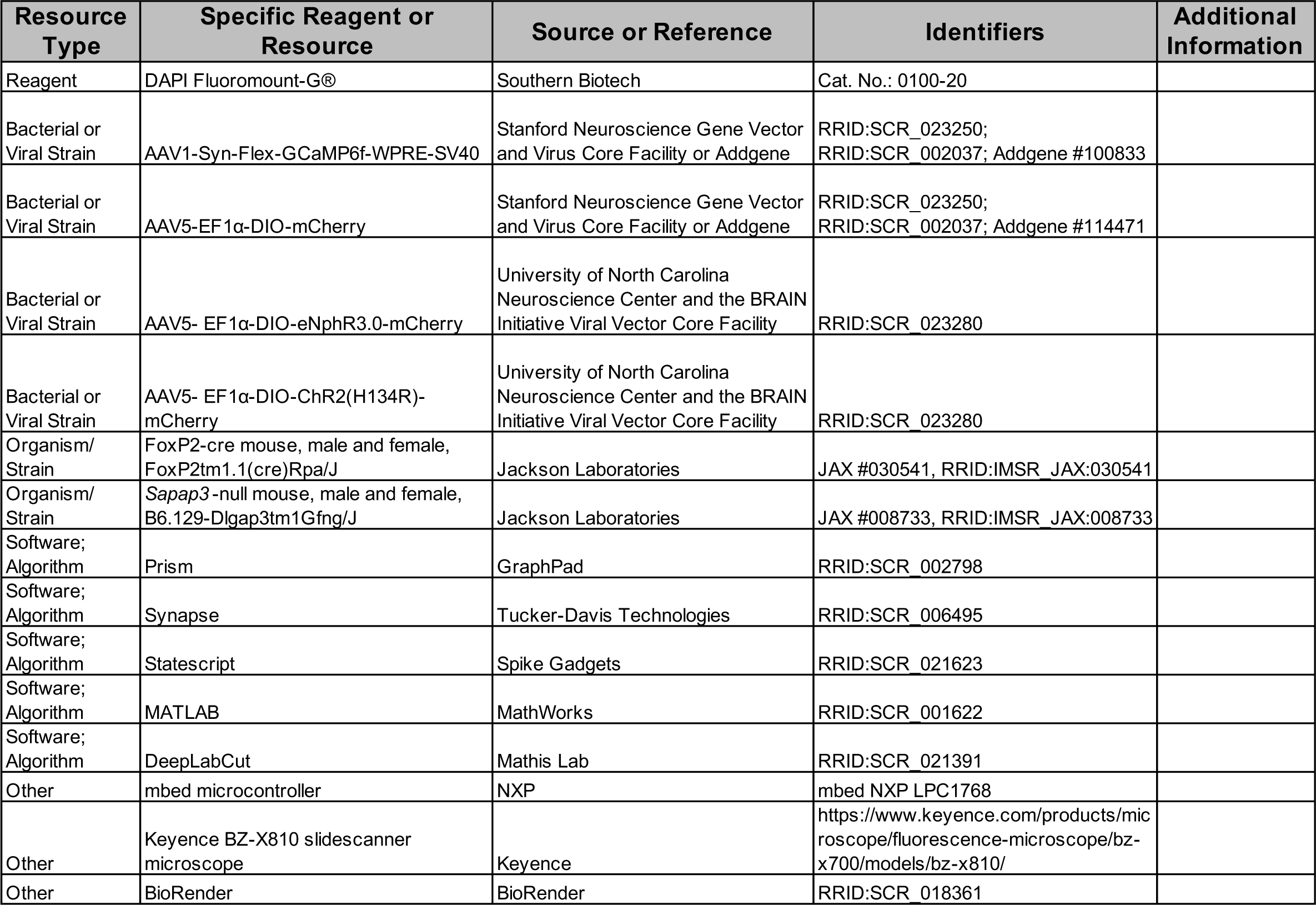

